# Neural Measures of Human Decision Making Track Evidence Accumulation in Learned Space

**DOI:** 10.64898/2026.01.22.701207

**Authors:** Arianna Thoksakis, Edward F. Ester

## Abstract

Neural decision-making flexibly integrates evidence across sensory, mnemonic, and semantic domains. Yet prior demonstrations have focused on evidence that is either directly available in stimuli, retrieved from established representations, or computed relative to fixed perceptual frameworks. A fundamental question remains: does neural evidence accumulation extend to decisions based on evidence that must be *computed* through learned representational transformations? Here we show it does. We recorded scalp EEG while participants classified continuously-oriented visual stimuli into discrete categories defined by an experimenter-imposed boundary. Category-level evidence was operationalized as category coherence, or the angular distance between each stimulus and the learned boundary. We predicted that if neural decision mechanisms are truly domain-general, the centro-parietal positivity (CPP)—a scalp EEG potential indexing evidence accumulation—should scale with category coherence, and individual differences in CPP sensitivity should correlate with computational drift rates. Both predictions were supported: CPP slopes increased monotonically with category coherence, and individual differences in CPP slopes correlated with individual differences in drift rates across participants. These findings reveal that the brain’s decision machinery treats evidence identically regardless of its representational origin---whether externally available, pre-existing, or computed through learned transformations.

**Significance Statement:** Evidence accumulation is a universal principle by which brains convert information into decisions. Prior work demonstrates this mechanism for sensory evidence, memory retrieval, and evidence computed relative to fixed perceptual axes, but a critical gap remains: does it extend to decisions based entirely on learned, arbitrary rules? We show that it does. Specifically, we demonstrate that the centro-parietal positivity (CPP)—a neural marker of evidence accumulation—tracks decisions in rule-defined category space, with buildup rates that scale with distance from learned boundaries and correlate with computational measures of evidence-accumulation rate. This reveals that the brain’s decision machinery is flexible, adapting to evidence in any representational framework, whether externally available or internally constructed.

---

A foundational framework in cognitive and systems neuroscience proposes that decisions arise from the continuous accumulation of evidence until a threshold is reached. Accumulation-to-threshold models successfully capture behavioral patterns across diverse domains, from simple perceptual discriminations to value-based choice (Ratcliff & McKoon, 2008). In these frameworks, the rate of accumulation governs both decision speed and accuracy, providing a computational bridge between stimulus information and choice behavior.

The centro-parietal positivity (CPP), a scalp EEG potential, is widely regarded as a neural signature of evidence accumulation (O’Connell et al., 2012; Kelly & O’Connell, 2013). The CPP is characterized by a gradual buildup of positive voltage that reaches an asymptote shortly before response execution. CPP buildup rate scales with evidence strength and predicts response time and accuracy. Recent work has extended CPP-linked accumulation to other domains. van Ede and Nobre (2024) showed that the CPP tracks decisions based on working memory representations, with buildup dynamics reflecting the time course of memory-guided deliberation. Tsvinev et al. (2025) demonstrated that CPP slope scales with semantic evidence in a manner analogous to sensory evidence. CPP dynamics have also been shown to predict subjective confidence independent of memory accuracy (Murphy et al., 2015; Dou et al., 2024). Thus, the CPP indexes evidence accumulation over multiple sources of information, including sensory input, confidence, working memory, and semantic knowledge.

Importantly, previous EEG studies have demonstrated that centroparietal signals can track decision-relevant evidence even when it is dissociated from the physical stimulus. Wyart et al. (2012) showed that during multi-sample orientation categorization, slow centroparietal EEG signals encoded each sample’s momentary “decision update”, i.e., its proximity to a fixed cardinal-diagonal category boundary — rather than its raw physical properties. Wyart et al. (2015) extended this finding to conditions of divided spatial attention, confirming that centroparietal decision signals are maintained even under attentional load. These studies established that centroparietal EEG activity can operate over transformed evidence variables. However, Wyart and colleagues used reverse correlation methods to isolate momentary decision-update signals from sequential stimulus samples, rather than measuring the CPP. Moreover, the category axis used in those studies (cardinal vs. diagonal) constitutes a fixed, universally shared perceptual distinction that remained constant across all participants and trials. Whether the CPP itself — as canonically defined and linked to accumulation-to-bound dynamics — tracks evidence defined by a boundary that is arbitrary, participant-specific, and newly acquired through learning has not been directly tested.

Categorization based on learned, arbitrary rules presents a different challenge: evidence is not directly available in the stimulus nor readily retrievable from existing knowledge structures. Instead, it must be computed by comparing sensory features to a newly learned, participant-specific framework. Whether the neural mechanisms underlying evidence accumulation can flexibly operate over evidence computed through such learned transformations remains unknown. This question has deep theoretical implications: If neural decision mechanisms are truly domain-general, they should track any evidence variable that governs behavior, regardless of its representational origin or whether it is computed through external sensing, memory retrieval, or learned transformations.

To test this hypothesis, we asked whether the CPP tracks decisions based on evidence computed in a learned category space. Specifically, we recorded scalp EEG while participants classified oriented visual stimuli into discrete categories defined by an arbitrary boundary, operationalizing category evidence as angular distance from the learned boundary. We predicted two outcomes: (1) if the CPP reflects domain-general accumulation, then CPP buildup rate should scale monotonically with category coherence; and (2) if category coherence parametrizes evidence strength analogously to sensory evidence metrics, then individual differences in CPP sensitivity should covary with computational measures of evidence-accumulation rate from drift-diffusion modeling. We report support for both predictions.

## Methods

### Data and Code Availability

Stimulus presentation software, EEG preprocessing routines, preprocessed EEG data, and analytic software sufficient to reproduce each manuscript figure can be found on the Open Sciences Framework: https://doi.org/10.17605/OSF.IO/QV8MA

### Sample Characteristics

A total of 41 human volunteers (both sexes) enrolled in this study. Each volunteer was recruited from the local university community, aged 18-40, self-reported normal or corrected-to-normal visual acuity and participated in a single 2.5-hour testing session in exchange for course credit or monetary remuneration ($15/h). We had no a priori reason to expect that task performance or EEG results would vary as a function of race, ethnicity, or any other immutable characteristic; therefore, to better protect volunteers’ privacy we did not collect this information.

### Data Exclusion

Data from one participant was discarded due to chance categorization performance (53%). Data from a second volunteer was discarded due to their voluntary withdrawal from the study. Data from a third volunteer was discarded due to experimenter error: the wrong EEG recording montage was selected and data from 32 of 64 scalp electrodes were irretrievably lost. Thus, the data reported here reflect the 38 remaining volunteers.

### Experimental Setup and Testing Environment

Stimuli were generated in MATLAB and rendered on a 27-inch g-sync enabled LCD monitor cycling at 240 Hz via Psychophysics Toolbox software extensions. The monitor resolution was fixed at 1920 × 1080 pixels. Participants were seated 85 cm from the display, with head position fixed by an SR Research chin rest. Under these conditions, 1 degree of visual angle (DVA) corresponds to 49.38 pixels. Stimuli in each experiment were apertures containing 350 iso-oriented bars with inner and outer radii of 2 and 9 DVA, respectively. Stimuli flickered at 30 Hz (50% duty cycle), and each bar in the aperture (1 DVA length, 8 pixel stroke width) was randomly replotted at the beginning of each “up” cycle to discourage volunteers from foveating individual bars. Participants reported category judgments by pressing keys on a standard USB-powered keyboard.

### Training Categorization Task

Volunteers learned to classify orientation stimuli into discrete groups according to an arbitrary (i.e., experimenter-chosen) boundary by completing a training task (e.g., Ester et al., 2020; Caron & Ester, 2025). Volunteers viewed apertures containing iso-oriented bars and reported whether they were members of “Category 1” or “Category 2” by pressing the “Z” or “?” key (respectively) (Figure 1A). Stimuli were presented for 3 seconds and followed by a 1.5-2.0 sec inter-trial interval (randomly and independently selected on each trial). Bars within the aperture were randomly replotted within the stimulus aperture at the beginning of each “up” cycle to discourage participants from foveating any one stimulus.

**Figure 1.**
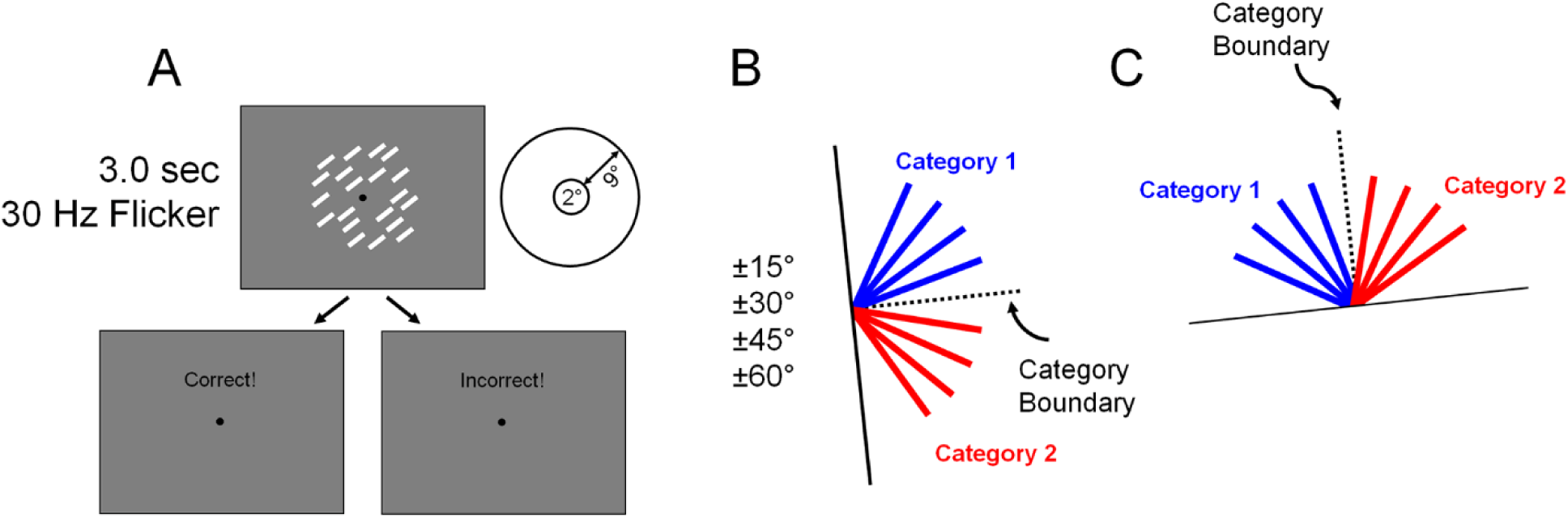
Training Task. (A) Trial schematic. Participants viewed displays of iso-oriented bars and classified them into discrete categories. Note that stimuli are not drawn to scale; see Training Categorization Task for specific parameters. (B-C) Examples showing the category boundary (black dashed line) and orientation values (±15°, ±30°, ±45°, ±60°). Each participant was assigned a unique and randomly selected category boundary on the interval [0, 180). The main categorization task was identical to the training task, with the exception that orientation values were drawn from the set ±2°, ±5°, ±15°, and ±45° relative to each participant’s category boundary.

Category membership was defined by a boundary whose orientation was randomly and independently selected for each participant on the interval [0°, 179°). We defined a set of stimuli tilted ±15°, ±30°, ±45°, and ±60° relative to each participant’s category boundary and assigned stimuli tilted counterclockwise to the boundary to Category 1 and stimuli tilted clockwise to the boundary to Category 2 (Figure 1B-C). Feedback – i.e., “correct!” vs. “incorrect!” was presented for 1 second at the end of each trial. Failures to respond within the 3-second stimulus window were treated as incorrect. Participants received verbal instructions describing key features of the task. They were told that there existed an (unknown) boundary that divided stimuli into different categories, and that their task was to identify this boundary through trial-and-error. In general, and with the benefit of these instructions, volunteers reached asymptotic performance by the 2^nd^ or 3^rd^ block, Figure 2). Participants then completed the main experiment using the same (bespoke) category boundary imposed during training. Each volunteer completed 4 blocks of 32 trials in the training task.

**Figure 2.**
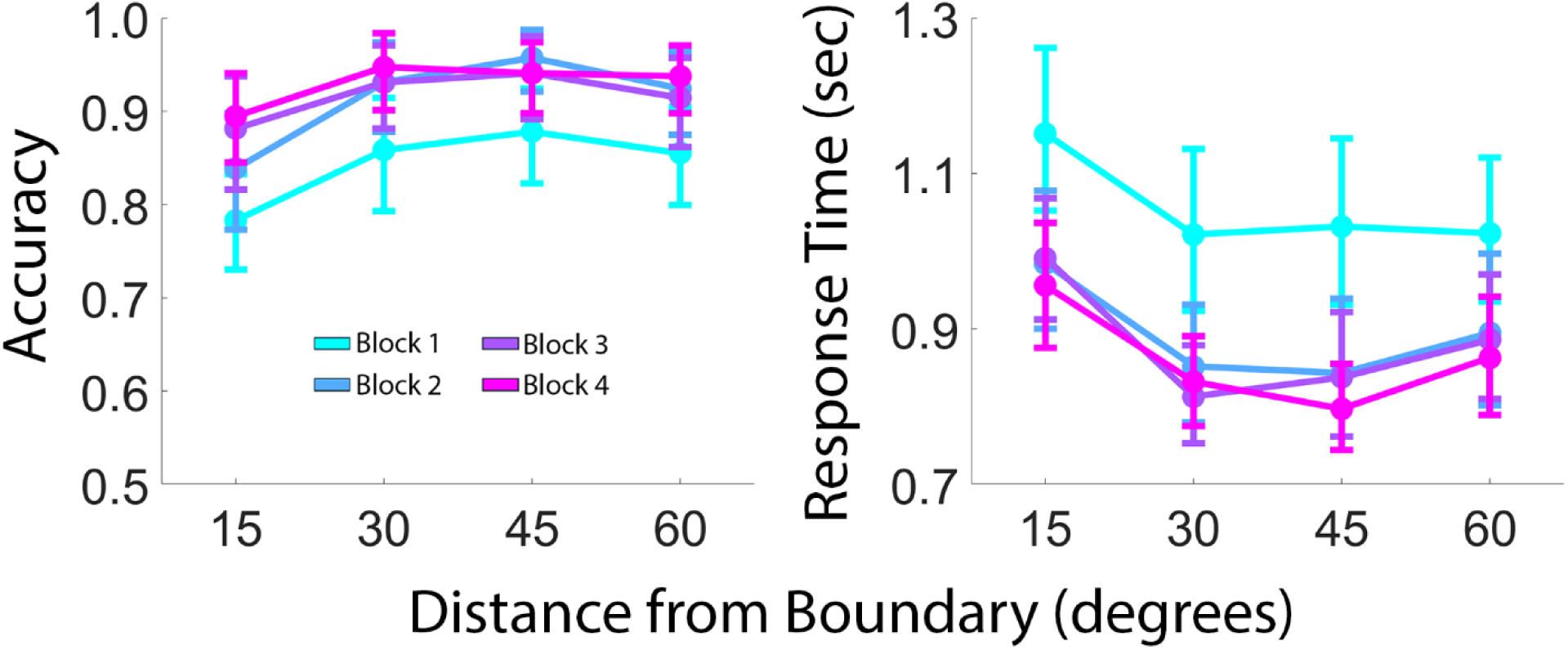
Training Task. Accuracy and Response Time are plotted as a function of block number (1-4). Participants learned the task rapidly, reaching asymptotic accuracy and response speed within 2–3 blocks. Error bars depict 95% confidence intervals.

### Main Categorization Task

After reaching proficiency on the training task, participants began the main experiment. Trial timing was identical to the training task. We defined a set of oriented stimuli ±2°, ±5°, ±15°, and ±45° relative to the participant’s category boundary, with negative values assigned to Category 1 and positive values assigned to Category 2. We included orientation values not presented during training (e.g., ±2° and ±5°) to make the task more difficult and to encourage participants to solve the task by applying category-defining rules vs. rote memorization. Participants reported the category membership of presented stimuli by pressing the “z” and “?” keys (for Categories 1 and 2, respectively) at any point during each three-second trial. Failures to respond within the 3 second trial window were treated as incorrect. We also included a small proportion of trials (11%) where the presented stimulus precisely matched the category boundary, i.e., ±0°. We did not analyze data from these trials in this study. Participants were not informed of this manipulation, and to maintain the ruse participants received randomized feedback at the end of these trials. Participants completed 5 (N = 1), 6 (N = 1), 7 (N = 1), 8 (N = 1), 12 (N = 1), 13 (N = 1), 14 (N = 1), 15 (N = 2), or 18 (N = 29) blocks of 36 trials as time permitted during a single 2.5-hour testing session.

### Behavioral Analyses and Drift Diffusion Model Fitting

Task performance was quantified by sorting participants’ accuracies and response times according to category coherence, or the angular distance separating to-be-classified stimuli from their category boundary, i.e., ±2°, ±5°, ±15°, ±45°. We observed no performance asymmetries across stimuli rotated anticlockwise and clockwise from participants’ boundaries (i.e., negative and positive orientation values, respectively), so we pooled and averaged the data across these conditions. We also excluded responses occurring < 0.4 sec relative to trial start, which we deemed implausibly fast. This resulted in an average (±1 S.E.M.) loss of 0.66% (±0.24%) of trials per participants.

To relate behavioral performance to evidence accumulation dynamics, a drift–diffusion model (DDM) was fit to each participant’s categorization data using the pyDDM framework (Heathcote et al., 2019; Shinn et al., 2020), which provides flexible tools for specifying and fitting stochastic accumulator models to choice–RT distributions. We fit the model to each participant’s data from the main categorization task, using accuracy and response time as joint constraints on the parameters. The model assumed a standard DDM architecture with three free parameters: drift rate *v*, decision bound *a*, and non-decision time *T*_*er*_. Drift rate governs the average rate at which noisy evidence is accumulated toward a decision boundary; the bound reflects the amount of evidence required to trigger a response; and non-decision time captures sensory encoding and motor execution processes that are assumed not to depend on the decision itself. For each participant, reaction times were expressed in seconds and coded with respect to the category choice (upper vs. lower decision boundary), and error and correct trials were both included. To capture the effects of category evidence, the drift rate parameter was allowed to vary as a function of category coherence. Note that category coherence was randomized over trials, and thus, participants had no way of knowing whether a given trial would contain small vs. large amounts of evidence. Therefore, we held the parameters for the decision bound and non-decision time constant across category coherence levels within participants. This parameterization implements the assumption that changes in difficulty primarily reflect changes in the strength of category-level evidence, rather than changes in response caution or peripheral processing. Model parameters were estimated by minimizing the discrepancy between the empirical and predicted joint distributions of response time and choice for each subject and condition using pyDDM’s built-in fitting routines.

To assess model fit quality, we compared the model’s predicted accuracy and mean RT against observed values for each coherence level and computed group-level R² values. We also generated quantile-probability plots by extracting the .1, .3, .5, .7, and .9 quantiles of the observed and predicted RT distributions separately for correct and error trials at each coherence level. To accommodate trials falling outside the model’s predicted RT range (e.g., contaminant responses), we included a 2% uniform mixture component in the model likelihood (Ratcliff & Tuerlinckx, 2002). Similarly, to evaluate whether a more flexible parameterization better accounted for the data, we compared the original model (M1; drift rate varies by coherence, boundary and non-decision time fixed) against two alternatives: a model in which drift rate and decision boundary both varied across coherence levels (M2), and a model in which drift rate, decision boundary, and non-decision time all varied across coherence levels (M3). Models were compared using Bayesian Information Criterion (BIC).

### EEG Recording and Preprocessing

We recorded continuous EEG from 63 scalp electrodes using a BrainProducts actiCHamp Plus system. Online recordings were referenced to an electrode placed over the left mastoid (10-10 site TP9) and digitized at 1 kHz. After recording, the following preprocessing steps were applied, in order: (1) resampling (from 1 kHz to 250 Hz), (2) high-pass filtering (zero-phase forward-and-reverse filters with cutoffs of 0.5 Hz), (3) detection and removal of noisy channels, (5) detection and reconstruction of noisy epochs using artifact subspace reconstruction, (6) interpolation of missing electrodes, (7) re-referencing to the mean voltage of all electrodes, (8) epoching (from -1 to +4 sec around stimulus onset), and (9) application of a surface Laplacian to minimize spatial cross-contamination across electrodes (Perrin et al., 1989). Steps 1-8 were implemented using EEGLAB functions (v. 2025.0 Delorme & Makeig, 2004); step 9 was implemented using custom software. In lieu of low-pass filtering, we smoothed stimulus- and response-locked ERP time courses with a 100 ms gaussian kernel. <u>CPP Analyses</u>. Following prior work (O’Connell et al., 2012; Twomey et al., 2016; van Ede & Nobre, 2024) we defined the centro-parietal positivity (CPP) as the average voltage measured over 10-20 electrode sites CP1, CPz, CP2, P1, Pz, and P2. The CPP buildup rate was defined as the slope of a line fit to each participant’s average CPP waveform over an interval spanning -1.0 to 0.0 sec relative to response onset. We used CPP slope rather than amplitude as our primary metric because slope is more directly related to the construct of evidence accumulation.

To test whether CPP slope dynamics reflected the same accumulation process captured by our drift-diffusion model, we correlated across-condition changes in drift rate with across-condition changes in CPP slope for each participant. For each participant, we computed a coherence-drift rate slope (reflecting the change in drift rate across the four coherence levels, Figure 3C) and a coherence-CPP slope (reflecting the change in CPP slope across the four coherence levels, Figure 4C). These across-condition slopes were then correlated across participants using Pearson correlation. This analysis tests whether individuals who show larger behavioral sensitivity to coherence (steeper drift-rate functions) also show correspondingly larger neural sensitivity to coherence (steeper CPP-slope functions).

**Figure 3.**
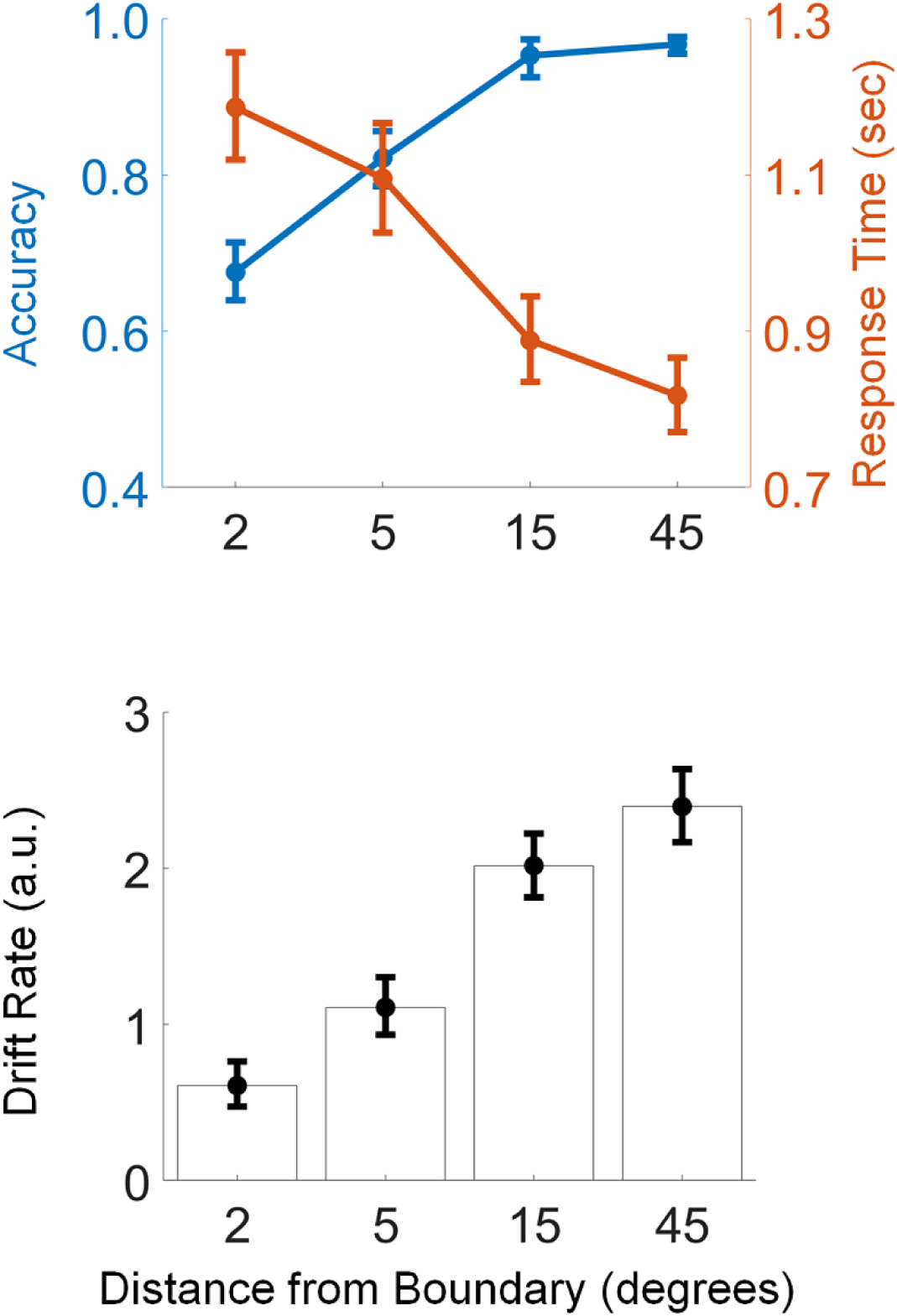
Category Coherence and Evidence Accumulation. (A) Accuracy increased monotonically with category coherence, as did response speed. (B) Drift rates from DDM fits scaled with coherence, indicating that evidence strength drove difficulty effects. Error bars depict 95% confidence intervals. a.u., arbitrary units.

**Figure 4.**
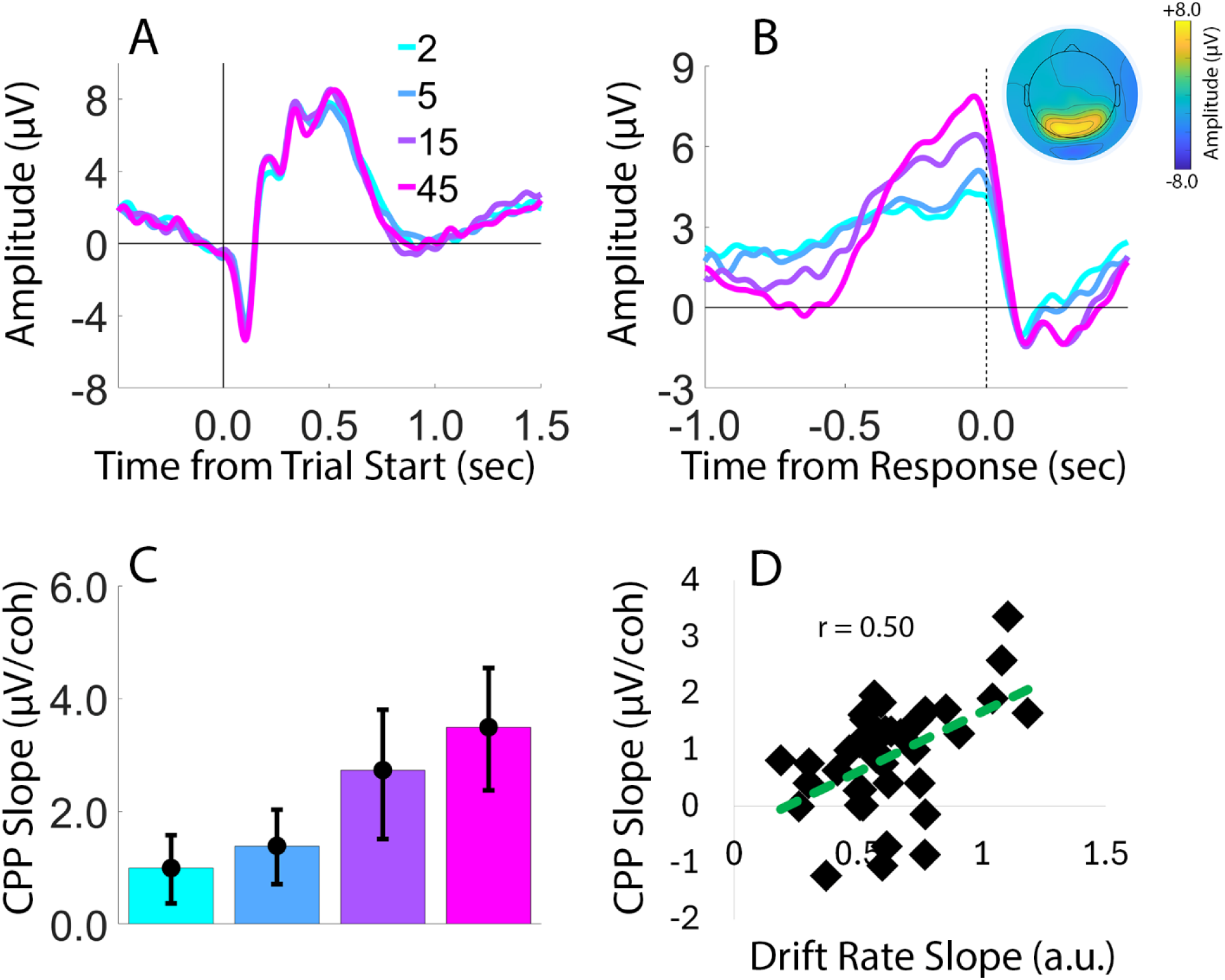
CPP Tracks Category Coherence and Aligns with Computational Evidence Accumulation. (A–B) Stimulus- and response-locked CPP waveforms. Response-locked CPP waveforms show ramping buildup preceding responses, with steeper slopes for high-coherence trials. Inset shows the distribution of voltages over scalp electrode sites, averaged from a period spanning - 300 to 0 ms prior to response. (C) CPP slopes increased monotonically with category coherence. (D) Across-participant correlation between drift-rate slope and CPP-slope. Error bars depict 95% confidence intervals. μV, microvolts. a.u., arbitrary units

### Lateralized Beta Power Analysis

To confirm that coherence-dependent modulations were specific to the CPP and were not mirrored in downstream effector-specific processes, we examined lateralized beta-band power (15–25 Hz) over motor cortex as a function of category coherence and response hand. Beta-band power was computed by applying a continuous wavelet transform to response-locked EEG data from frontocentral electrode site pairs C3/4 and FC3/4 For each participant and coherence level, we computed a lateralized motor index as the difference between contralateral and ipsilateral power relative to the responding hand (contralateral minus ipsilateral), which quantifies effector-specific motor preparation. Response-locked motor beta waveforms were then time-locked to the response and averaged across trials for each coherence level, yielding four beta waveforms per participant. To assess the buildup rate of lateralized motor activity, we fit a line to each participant’s average response-locked beta waveform over the interval from −1.0 to 0.0 s relative to response onset, mirroring the approach used for CPP analysis. Motor beta slopes were then compared across coherence levels using a one-way repeated-measures ANOVA to determine whether effector preparation varied as a function of category evidence strength.

### Statistical Analyses

Unless stated otherwise, all statistical tests were based on standard approaches (t-tests, ANOVA). Where null results were theoretically relevant, we supplemented frequentist tests with Bayesian analyses to quantify evidence for the null hypothesis. Bayes factors were computed using the bayesFactor MATLAB toolbox (Krekelberg, 2024), which implements the default JZS priors described by Rouder et al. (2009, 2012). We interpret Bayes factors using the conventions of Jeffreys (1961), with BF₀₁ > 5 indicating substantial evidence for the null.

## Results

### Behavioral performance and diffusion modeling

Participants learned the arbitrary category boundary quickly during the training task. Accuracy increased and response times decreased over early blocks as a function of category coherence, with no interaction between these factors (accuracy: main effect of task block, *F*(3,111) = 10.583, *p* < 10^−4^, *η*^2^ = 0.222; main effect of category coherence, *F*(3,111) = 10.502, *p* < 10^−4^, *η*^2^ = 0.221; interaction, *F*(9,333) < 1, *p* = 0.535; RT: main effect of block, *F*(3,111) = 23.634, *p* < 10^−4^, *η*^2^ = 0.387; main effect of category coherence, *F*(3,111) = 21.671, *p* < 10^−4^, *η*^2^ = 0.369; interaction, *F*(9,333) < 1, *p* = 0.685; Figure 2). This pattern indicates rapid acquisition of the categorization rule without systematic asymmetries between categories.

In the main categorization task, accuracy increased and response times decreased monotonically with the category coherence (Figure 3A). A one-way repeated-measures ANOVA confirmed a strong main effect of category coherence on accuracy (*F*(3,111) = 190.81, *p* < 10^−4^, *η*^2^ = 0.837) and on response times (*F*(3,111) = 126.72, *p* < 10^−4^, *η*^2^ = 0.774), with better and faster performance for stimuli distal from the boundary.

To relate behavioral performance during the categorization task to evidence-accumulation dynamics, a drift–diffusion model was fit to each participant’s accuracy and RT distributions, allowing drift rate *v* to vary with category coherence while holding the decision bound *a* and non-decision time *T*_*er*_ fixed across coherence levels. Drift rates increased monotonically with coherence (one-way repeated-measures ANOVA: *F*(3,111) = 219.98, *p* < 10^−5^, *η*^2^ = 0.85; Figure 3B), consistent with a positive relationship between category coherence and evidence accumulation rates. The model provided an excellent fit to the behavioral data. Group-level predicted and observed accuracy (R² = 0.997) and mean RT (R² = 0.993) were in close correspondence across all four coherence levels. Quantile-probability plots confirmed that the model captured the shape of correct and error RT distributions, with only modest deviations at the upper quantiles of error trials where trial counts were low (Extended Data Figure 3-1).

To evaluate whether a more flexible parameterization better accounted for the data, we compared the original model (M1; drift rate varies by coherence, boundary and non-decision time fixed) against two alternatives: a model in which drift rate and decision boundary both varied across coherence levels (M2), and a model in which drift rate, decision boundary, and non-decision time all varied across coherence levels (M3). Models were compared using BIC. M1 was favored in 34 of 38 participants, M2 in 3 of 38 participants, and M3 in 1 of 38 participants (mean ΔBIC: M1 − M2 = -11.2; M1 − M3 = -16.7), confirming that the constrained parameterization provides the best account of the data and that difficulty effects in this design are driven primarily by changes in evidence strength.

### CPP Buildup tracks category coherence

CPP activity was quantified as the average voltage over electrodes CP1, CPz, CP2, P1, Pz, and P2, time-locked to trial onset and to the response (Figure 3A–B). Response-locked CPP waveforms exhibited a ramping buildup preceding button presses, and this buildup was visibly steeper for stimuli farther from the category boundary than for boundary-adjacent stimuli. CPP buildup rate was operationalized as the slope of a line fit to each condition’s average response-locked CPP over the interval from −1.0 to 0.0 s relative to response onset (Figure 4B–C). A one-way repeated-measures ANOVA on these slopes revealed a significant main effect of category coherence (*F*(3,111) = 22.53, *p* < 10^−4^, *η*^2^ = 0.378), such that CPP slopes increased with category coherence (Figure 4C), consistent with faster neural accumulation for high-coherence category decisions.

### CPP slopes covary with behavioral drift rates

To test whether the CPP reflects the same accumulation process captured by the diffusion model, across-condition changes in drift rate were compared to across-condition changes in CPP slope for each participant. For each participant, a coherence–drift rate slope (from Figure 3C) and a coherence–CPP slope (from Figure 4C) were computed, and these across-condition slopes were then correlated across participants (Figure 4D). Participants who showed larger increases in drift rate across coherence levels also showed larger increases in CPP slope, yielding a significant positive correlation (*r* = 0.50, *p* < 1*e* − 04; 95% CI [0.21, 0.71]; Figure 4D). This coupling between model-derived drift and neural buildup supports the interpretation of the CPP as a neural expression of the decision variable governing category judgments, rather than a generic evoked response. Notably, because drift rates are estimated entirely from behavioral data, this between-subject correlation is not easily explained by condition-level confounds such as differential overlap of stimulus-evoked potentials with the response-locked measurement window (see Discussion).

### Motor activity does not account for CPP Effects

Finally, to rule out the possibility that CPP modulations reflected differential motor preparation, lateralized beta-band power (15-25 Hz) over motor cortex was examined as a function of response hand and category coherence. Beta power was computed over frontocentral electrodes and expressed as a normalized contralateral–ipsilateral difference relative to the responding hand, then sorted by category coherence (Figure 5). For each participant, we fit slopes to the response-locked beta waveforms over the interval spanning -1000 to 0 ms relative to response onset, one slope per coherence level. As expected, contralateral beta power decreased relative to ipsilateral power prior to responses, but the slope of this decrease did not vary systematically with category coherence [F(3,111) = 1.27, p = 0.28, η^2^ = 0.03]. A Bayesian repeated-measures ANOVA yielded very strong evidence for the null hypothesis (BF_01_ = 36.76), confirming that effector-specific motor preparation was not modulated by category coherence. The absence of significant coherence effects on motor beta slopes, together with robust coherence and drift-rate dependencies of the CPP, is consistent with the interpretation that the CPP effects reflect central evidence-accumulation processes rather than downstream motor activity.

**Figure 5.**
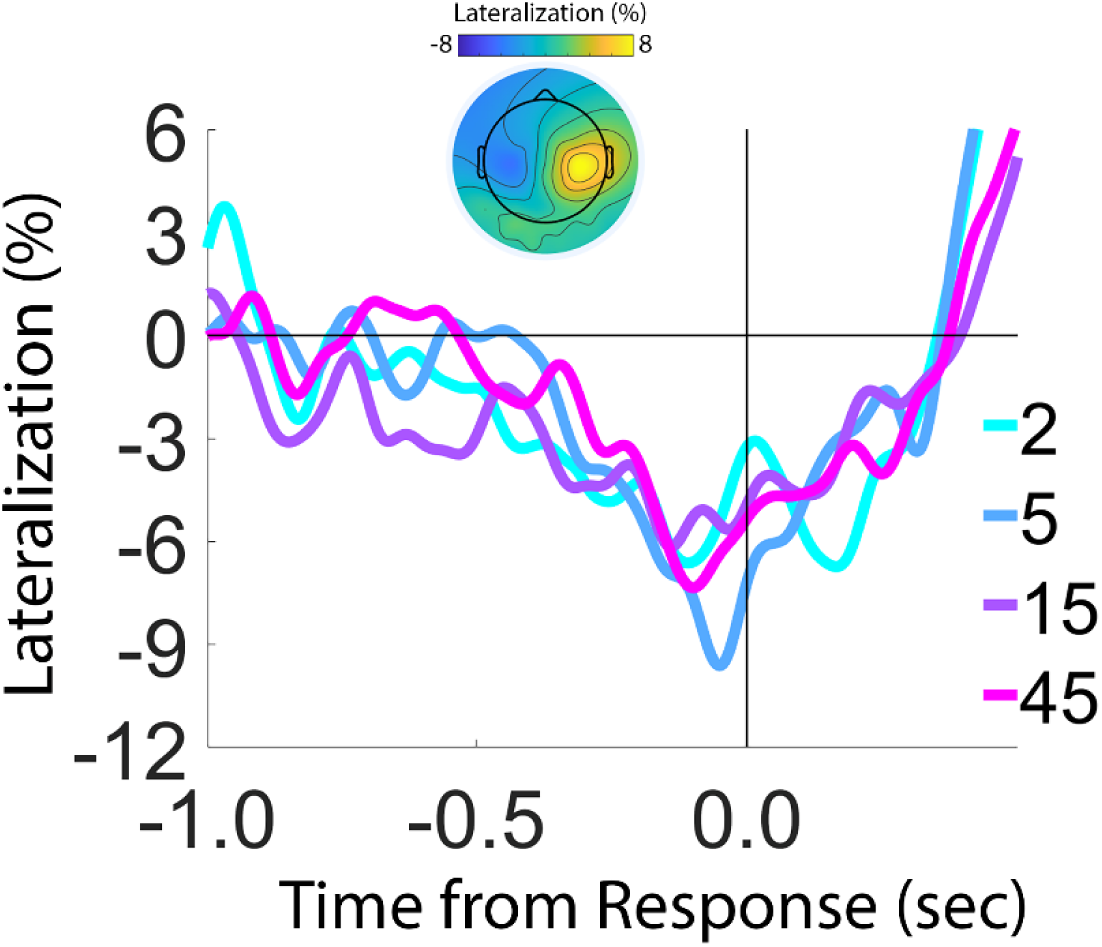
Motor Preparation Does Not Account for CPP Effects. Lateralized motor beta power as a function of category coherence. Although beta power showed the expected contralateral suppression prior to response, it was insensitive to category coherence (BF_01_ = 36.76), indicating that CPP modulations reflect central decision processes rather than downstream motor preparation. Inset shows the scalp distribution of lateralized beta power.

## Discussion

Evidence accumulation is a central principle by which neural systems convert information into decisions. A foundational assumption in decision neuroscience is that this accumulation process should be domain-general—flexibly operating over any decision variable that governs behavior, regardless of its representational source. Previous work has shown that the centro-parietal positivity (CPP), a neural index of evidence accumulation, tracks decisions based on sensory input, working memory, semantic knowledge, and subjective confidence (O’Connell et al., 2012; Kelly & O’Connell, 2013; Murphy et al., 2015; van Ede & Nobre, 2024; Tsvinev et al., 2025; Dou et al., 2024). Other studies have demonstrated that centroparietal EEG signals track decision-relevant evidence even when it must be computed by transforming physical stimulus properties relative to a fixed category axis (Wyart et al., 2012, 2015). However, whether the CPP extends to decisions based on evidence computed through newly learned, arbitrary representational transformations has remained untested. Here, we addressed this question directly.

Our findings strongly support domain-generality. Two key results converge on this conclusion. First, CPP buildup rate scaled monotonically with category evidence (angular distance from a learned decision boundary), demonstrating that neural accumulation operates identically over evidence computed through learned rules. Second, individual differences in CPP sensitivity to this computed evidence covaried with individual differences in drift-rate sensitivity from computational modeling, mirroring the CPP–DDM relationship established for sensory evidence. Critically, this coupling indicates that the CPP indexes the same underlying decision variable across vastly different representational formats—sensory, mnemonic, semantic, and now computed-through-learning.

A key distinction concerns the nature and origin of the evidence being accumulated. In standard perceptual paradigms, evidence is typically stimulus-defined and manipulated via sensory noise (e.g., the coherence of a moving dot stimulus). In semantic tasks, evidence is retrieved from long-term memory (e.g., knowledge of U.S. state populations; Tsvinev et al., 2025). In orientation categorization paradigms, evidence can be extracted by computing the distance between a physical stimulus and a category axis (Wyart et al., 2012, 2015). However, in those studies, the category axis (cardinal vs. diagonal) constituted a fixed, perceptually salient distinction shared across all participants, and evidence was extracted from sequential stimulus samples using reverse correlation — an approach that isolates momentary decision-update signals rather than the sustained, response-locked accumulation dynamics characteristic of the CPP (O’Connell et al., 2012). By contrast, in the present design, evidence was computed on each trial by applying a learned transformation: participants mapped sensory features (orientation) onto boundary-centered decision coordinates (angular distance from the learned boundary). The relevant decision variable—distance-to-boundary—does not exist in the stimulus itself but must be calculated by comparing the sensory input to an internally represented, idiosyncratic category boundary. Unlike the fixed cardinal-diagonal axis used in prior work, this boundary was arbitrary, participant-specific, and acquired through trial-and-error learning during the experimental session. The same physical orientation can therefore convey different amounts of evidence for different observers, depending on their learned boundary. Critically, these results reveal that CPP-linked accumulation is not tethered to sensory evidence sampled directly from the environment or to evidence computed relative to fixed perceptual frameworks but instead reflects integration of task-relevant evidence regardless of its computational origin. This functional flexibility suggests that neural decision mechanisms reflected by the CPP can adaptively incorporate information from circuits that perform task-specific transformations, whether those transformations involve sensory filtering, memory retrieval, or rule application.

A critical finding is the individual-differences correlation between CPP-slope sensitivity and drift-rate sensitivity. This coupling is notable for two reasons. First, it replicates the established CPP–DDM relationship from perceptual decision-making (Kelly et al., 2021), extending it to evidence computed through learned transformations. Second, and more fundamentally, it demonstrates that the CPP and drift rate index the same underlying decision *process* across representational domains. The brain’s neural machinery for accumulating sensory evidence and its neural machinery for accumulating computed evidence are not separate systems—they are a single, flexible mechanism organized around decision variables.

The observed CPP modulations are unlikely to reflect motor preparation. To confirm that coherence effects were specific to the CPP — an abstract, effector-independent decision signal — and were not mirrored in downstream effector-specific processes, we examined lateralized beta-band activity over motor cortex. Although beta power exhibited the expected contralateral suppression prior to response, the slope of lateralized beta buildup did not significantly differ across coherence levels, suggesting that effector preparation was not detectably modulated by category coherence. We note that several previous studies have reported difficulty-dependent motor beta buildup rates in perceptual decision tasks (De Lange et al., 2013; Steinemann et al., 2018). One possible explanation for the difference is that our categorization task may involve an additional mapping step — from accumulated category evidence to a motor response — that partially decouples motor preparation from the graded evidence signal. Nevertheless, the CPP itself scaled systematically with category evidence and covaried with drift rate, and prior work has consistently dissociated CPP dynamics from motor preparation, demonstrating that CPP modulations generalize across response effectors (O’Connell et al., 2012; Twomey et al., 2016). Together, these findings support the interpretation of the CPP as a central decision variable operating over category-level evidence, rather than a byproduct of downstream motor planning.

Our findings establish a critical link between human electrophysiological evidence and monkey single-unit neurophysiology. In lateral intraparietal area (LIP) of macaques, neurons show ramping activity during perceptual decision-making that reflects evidence accumulation toward a decision bound (Roitman & Shadlen, 2002; Shadlen & Newsome, 2001). Importantly, these same LIP neurons also encode learned category membership for visual stimuli, such as motion direction (Freedman & Assad, 2006; Swaminathan & Freedman, 2012). Moreover, LIP neurons exhibit ramping dynamics during memory-based categorical decisions, where evidence is sampled sequentially from internal memory representations when sensory stimulus is no longer present (Shushruth et al., 2022). Reversible inactivation of LIP impairs categorization performance in multiple task contexts (Zhou & Freedman, 2019; Zhou et al., 2023; Peysakhovich et al., 2024), demonstrating a causal role for LIP in visual categorical decisions. Freedman and Assad (2011) proposed that categorization and perceptual decision-making may engage a common neural mechanism in parietal cortex, where evidence is accumulated regardless of whether it derives from sensory properties or learned categorical structure. Our results provide the human analog of this proposal: the CPP, thought to reflect parietal accumulation processes homologous to those in monkey LIP, tracks evidence defined in learned category space. This cross-species convergence strengthens the interpretation that parietal cortex implements domain-general evidence accumulation that operates flexibly across representational formats, whether sensory features, semantic knowledge, or task-constructed categorical structure.

One methodological consideration is that the sudden onset of each visual stimulus elicits a sensory-evoked response in the ERP that varies in its temporal overlap with the pre-response measurement window as a function of response time and, consequently, category coherence. Several approaches have been developed to minimize this concern, including stimulus lead-in periods that temporally separate sensory onset from evidence onset (Kelly et al., 2021) and deconvolution techniques such as RIDE that decompose stimulus-locked and response-locked ERP components (Steinemann et al., 2018; Grogan et al., 2023). The present design did not incorporate these controls. Nevertheless, the across-participant correlation between CPP-slope sensitivity and drift-rate sensitivity (Figure 4D) reflects between-subject covariation in the functional coupling between neural buildup and computational evidence strength. This pattern is not straightforwardly predicted by differential sensory contamination, which would need to systematically track individual differences in drift-rate sensitivity rather than merely condition-level RT differences. Future work incorporating stimulus lead-in periods or deconvolution methods would provide further leverage on this issue.

Several additional limitations suggest directions for future work. Although our design provides strong evidence that the CPP reflects accumulation in learned category space, it does not isolate the locus of the transformation from sensory features to category evidence. Additionally, the task relied on a single, one-dimensional stimulus feature (orientation) and a deterministic category boundary. Extending this approach to multidimensional category spaces, probabilistic boundaries, or categories requiring integration across multiple features would clarify whether CPP-linked accumulation generalizes broadly across categorization regimes. Finally, categories in natural environments are often structured hierarchically or probabilistically, unlike the deterministic boundary used here (Ashby & Maddox, 2005). Testing whether CPP dynamics similarly track graded category evidence in more complex and ecologically rich categorization tasks would further establish the generality of these findings.

In summary, CPP buildup rate tracked evidence accumulation in a learned, boundary-centered representational space and covaried with drift-rate sensitivity across participants. These findings demonstrate that the CPP reflects integration of evidence computed through learned transformations, rather than being limited to sensory evidence sampled from the environment, fixed perceptual axes, or semantic knowledge retrieved from memory. The CPP thus indexes a flexible accumulation mechanism that adapts to whatever evidence coordinates are defined by task structure and acquired through learning. By showing that neural accumulation mechanisms operate over evidence defined in learned category space, the present results establish the representational flexibility of decision signals across perceptual, mnemonic, semantic, and categorical domains, and highlight a domain-general accumulation process capable of integrating diverse evidence representations, whether externally available, pre-existing, or constructed through learning and task demands.

## Author Contributions (CRediT Taxonomy)

Conceptualization: EFE, AT

Data Curation: AT, EFE

Formal Analysis: EFE, AT

Funding Acquisition: EFE

Investigation: AT, EFE

Methodology: AT, EFE

Project Administration: AT, EFE

Resources: EFE

Software: EFE, AT

Supervision: EFE

Validation: AT, EFE

Visualization: AT, EFE

Writing – Original Draft: EFE, AT

Writing – Review & Editing: EFE, AT

The authors declare no conflicts.

